# A *C. elegans* model for functional analysis of ADPKD variants in cilia, extracellular vesicles, and sensory signaling

**DOI:** 10.64898/2026.04.14.718433

**Authors:** Juan Wang, Carlos Nava Cruz, Jonathon D Walsh, Elizabeth desRanleau, Inna A Nikonorova, Maureen M Barr

## Abstract

Interpreting the pathogenic significance of missense variants in human disease gene candidates remains a major challenge in precision medicine. Autosomal dominant polycystic kidney disease (ADPKD) is the most common genetic cause of kidney failure and caused by mutations in the PKD1 or PKD2 genes that encode polycystin-1 and polycystin-2. Here, we establish *C. elegans* as a platform for the functional classification of PC2 variants by characterizing PKD-2^C180S^, the *C. elegans* ortholog of the likely pathogenic human variant PC2^C331S^. Using CRISPR/Cas9 endogenous genome editing combined with dual-color fluorescent reporters and super-resolution imaging, we show that PKD-2^C180S^ severely reduces protein stability, abolishes ciliary and extracellular vesicle (EV) localization, and eliminates sensory function comparable to a *pkd-2* null allele. In heterozygous animals, PKD-2^C180S^ is recessive and exerts no dominant-negative effect on wild-type PKD-2 trafficking, protein levels, or function, establishing that PKD-2 is haplosufficient in this model. PKD-2^C180S^ also abolishes ciliary and EV localization of the PC1 homolog LOV-1 and reduces LOV-1 cell body levels comparable to *pkd-2* null animals, consistent with PC2 functioning as a molecular chaperone for PC1 stability and trafficking. Genetic epistasis experiments show that PKD-2^C180S^ protein levels are unaffected in *lov-1* mutants, indicating that the PKD-2^C180S^ mutation acts prior to complex assembly. Quantitative analysis reveals that LOV-1•PKD-2 complexes are more stable at the ciliary membrane and more efficiently packaged into EVs than PKD-2 lacking LOV-1. Together, this work demonstrates that PC2^C331S^ may act recessively via loss of polycystin complex function and establishes a *C. elegans* pipeline for the mechanistic classification of ADPKD-associated variants.

## Introduction

Understanding the molecular and cellular mechanisms by which genetic variants contribute to human disease is a central challenge of the precision genomic medicine era (1). Autosomal dominant polycystic kidney disease (ADPKD) is a life-threatening and the most prevalent monogenic kidney disease and exemplifies the difficulties inherent in variant interpretation. Mutations in two genes, PKD1 and PKD2, account for approximately 95% of ADPKD cases (2, 3). PKD1 encodes polycystin-1 (PC1), a large 11-transmembrane-domain protein that combines structural features of adhesion GPCRs and transient receptor potential polycystin (TRPP) channels (4, 5), while PKD2 encodes polycystin-2 (PC2), a nonselective cation TRPP channel (6, 7). The scale of clinical variant data in these genes is substantial: ClinVar currently catalogs 5,006 and 1,037 variants in PKD1 and PKD2, respectively (“PKD1“[GENE] - ClinVar - NCBI; “PKD2“[GENE] - ClinVar - NCBI, accessed February 2026), underscoring the pressing need for systematic functional validation of ADPKD-associated variants.

Polycystins localize to cilia and contribute to signaling networks that constrain cyst initiation and growth (8–10). Reduced ciliary localization of the polycystin complex compromises polycystin-dependent signaling and represents a central disease mechanism in ADPKD (9–11). PC1 and PC2 have also been detected in urinary extracellular vesicles (EVs; (12–15)), which are emerging as drivers of cystogenesis and potential disease biomarkers (16). Thus, defects in polycystin ciliary and EV localization may represent a major class of pathogenesis mechanisms for ADPKD-associated variants. Modeling how missense mutations affect polycystin ciliary and EV localization and/or polycystin signaling provides a framework of functional assays to categorize variant effects and offers mechanistic insights into polycystin biology.

In this work, we establish *C. elegans* as a model to study how a likely pathogenic missense mutation, PC2 cysteine-331-to-serine (PC2^C331S^), affects ciliary and EV localization of the orthologous mutation PKD-2^C180S^. The *C. elegans* PC1 and PC2 homologs, LOV-1 and PKD-2, localize to male-specific sensory cilia, EVs, and are required for male mating behavior (17, 18). Male mating behavior assays provide a quantitative readout of polycystin function in living animals. *C. elegans* is currently the only *in vivo* model for observing polycystin localization to cilia and ciliary EVs in real time. Taking advantage of CRISPR/Cas9 genome editing, we studied the effects of PKD-2^C180S^, the mutation orthologous to PC2^C331S^, and its interaction with the PC1 homolog LOV-1 in an endogenous expression context.

Here, we demonstrate that *C. elegans* PKD-2^C180S^ abolishes ciliary and EV localization and disrupts polycystin-mediated sensory functions required for male mating behavior. We show that PKD-2^C180S^ is recessive for both localization and function, and acts as a loss-of-function allele that phenocopies the *pkd-2* null mutation. Importantly, PKD-2^C180S^ is unable to interact with wild-type PKD-2 or LOV-1, preventing PKD-2^C180S^ trafficking to cilia and ciliary EVs while also disrupting LOV-1 localization and stability. These findings reveal a genetic and cellular mechanism by which the PC2^C331S^ mutation acts and establish the utility of *C. elegans* as a platform for investigating the potential pathogenicity of patient-derived PC2 variants.

## Results

### Sensory functions and cellular mechanisms of the polycystin-2 C331S (PC2 ^C331S^) mutation in *C. elegans*

To establish *C. elegans* as a model for studying missense variants in PC2, we compared the AlphaFold-predicted *C. elegans* PKD-2 structure with the cryo-EM structure of human PC2 (PDB 6A70; (19)). The *C. elegans* PKD-2 AlphaFold-predicted structure aligned closely with the PC2 structure (Fig. 1A, Movie S1). In human PC2, cysteine 331 is located within the extracellular tetragonal opening of polycystin (TOP) domain, where C331 forms a disulfide bond with C444 to constrain the Finger 1 extension that locks into an adjacent subunit, thereby stabilizing the polycystin tetramer (20–23). Primary sequence alignment showed that the *C. elegans* TOP domain shares 40% identity with human PC2, with PKD-2 C180 corresponding to PC2 C331 (Fig. 1A-B, Fig. S1A-D, Movie S1).

**Figure 1.**
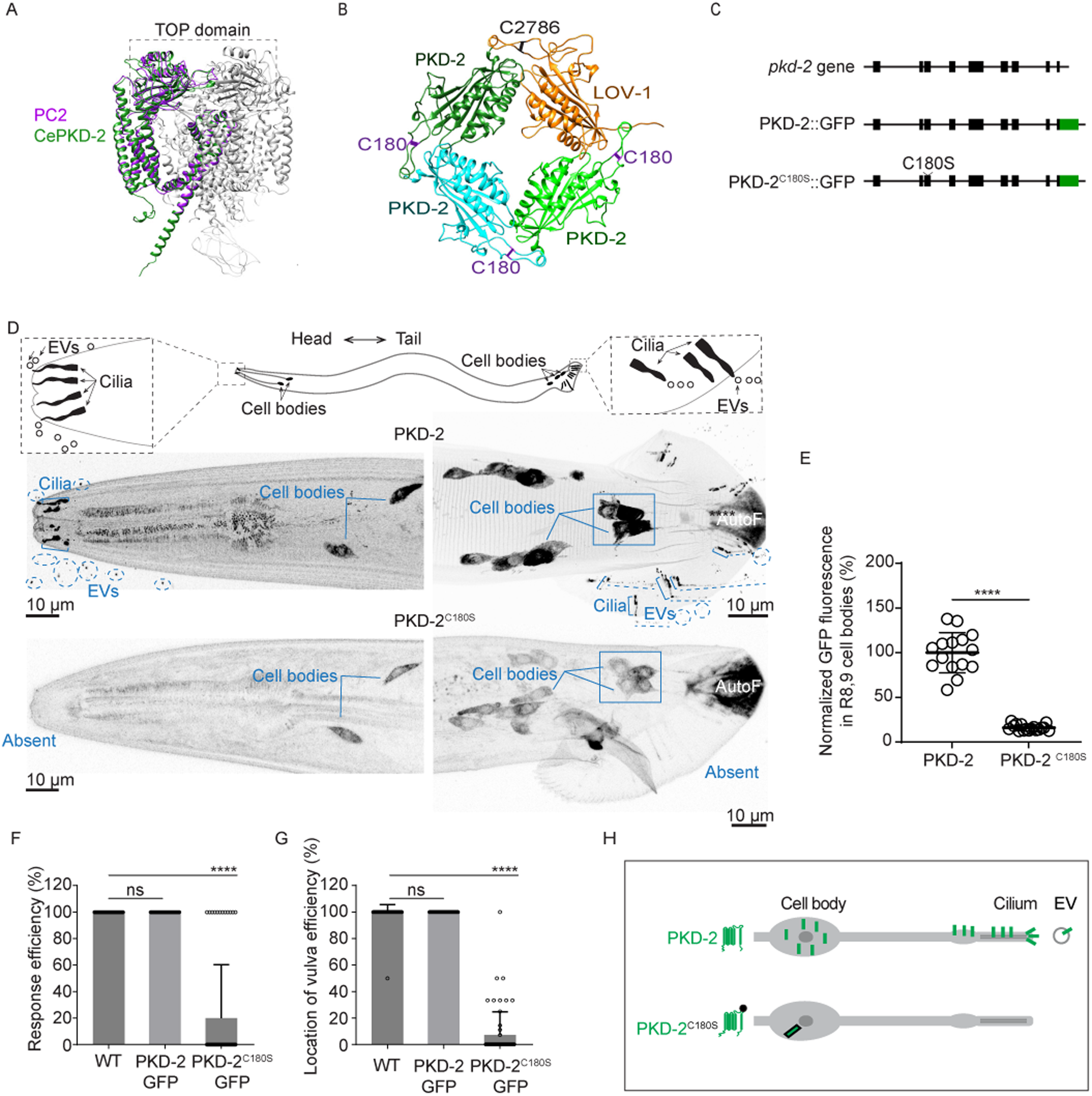
Modeling the orthologous PC2^C331S^ mutation in *C. elegans* rendered the mutated PKD-2^C180S^ protein functionally null and abolished both ciliary and EV localization. (A) Structural comparison of AlphaFold-predicted structures of CePKD-2 (green, UniProt: Q9U1S7, residues 60-619) and human PC2 (purple, residues 185-723, PDB: 6A70) in complex with PC1. (B) AlphaFold-predicted interactions between TOP domains of three CePKD-2 subunits and LOV-1, showing the disulfide bond at the interface of adjacent subunits. (C) Gene structure of *pkd-2* and genome editing strategy. *pkd-2(my121)* [PKD-2::GFP] is a CRISPR/Cas9-edited GFP knock-in allele inserted before the stop codon (24) *pkd-2(my154)* [PKD-2^C180S^::GFP] is a CRISPR/Cas9-edited allele in the *my121* background, changing cysteine180 to serine to visualize the effect of the PKD-2^C180S^ mutation on protein localization (see Methods). (D) The PKD-2^C180S^ mutation altered PKD-2 protein levels and localization. The cartoon depicts male-specific head and tail neurons expressing the *pkd-2* gene. The male tail has nine pairs of ray neurons, which are important for the mating behavior of contact response. Inset cartoons illustrate that PKD-2::GFP is enriched in cilia and released in EVs from ciliary tips into the environment. Representative micrographs show PKD-2::GFP enrichment in the cell bodies, sensory cilia, and EVs in wild-type animals (upper panels), whereas PKD-2^C180S^::GFP displays a weak signal in the cell bodies and is absent from sensory cilia and EVs (lower panels). Rectangles indicate neuronal cell bodies used for fluorescence intensity quantification in panel E. The posterior most portion of the male tail is autofluorescent (AutoF). Scale bars: 10 μm. (E) Quantification of PKD-2::GFP fluorescence intensity in the cell bodies of ray 8B and 9B neurons, as indicated by the rectangles in panel (D). The PKD-2^C180S^ mutation reduced protein levels to approximately 15% of those in wild-type animals. Data represent mean ± SD from at least three independent imaging sessions (n = 15 animals for PKD-2::GFP; n = 17 for PKD-2^C180S^::GFP). Statistical significance was determined by the Mann-Whitney test. **** indicates *p* < 0.0001. (F-G) Male mating behavior assays compared *pkd-2 (my154)* PKD-2^C180S^::GFP to wild-type *pkd-2* (+) and *pkd-2 (my121)* PKD-2::GFP. The PKD-2::GFP tagged allele *pkd-2 (my121)* males showed wild-type level mating efficiency in both the response to hermaphrodites (F) and vulva-location behavior (G). In contrast, *pkd-2 (my154)* PKD-2^C180S^::GFP males showed significantly reduced mating behavior efficiency compared to wild-type animals. Data represent mean ± SD from three independent blinded behavioral assays with 20 animals per genotype per assay. Statistical analysis was performed by Kruskal-Wallis test with Dunn’s multiple comparisons. **** indicates *p* < 0.0001; ns indicates *p* > 0.9999. (H) Summary: In wild-type sensory neurons, PKD-2 is enriched in cell bodies, cilia, and EVs. In contrast, PKD-2^C180S^ levels are reduced in cell bodies and PKD-2^C180S^ is not detected in cilia or EVs.

To track the molecular and cellular consequences of the PKD-2^C180S^ mutation in its native genomic context, we introduced the PKD-2^C180S^ mutation into a previously generated PKD-2::GFP strain (*pkd-2 (my121))* (24) through CRISPR/Cas9 editing at the *pkd-2* locus (Fig. 1C). This approach enabled us to track protein dynamics using super-resolution imaging in real time and *in vivo*.

In wild-type animals, PKD-2::GFP is enriched in cell bodies and is transported along dendrites to accumulate in male-specific sensory cilia located at distal dendrite tips (25, 26) PKD-2 is also shed in ciliary extracellular vesicles (EVs; Fig. 1D; (18, 24)). The PKD-2^C180S^ mutation completely abolished both ciliary and EV localization (Fig. 1D). Additionally, PKD-2^C180S^ reduced protein levels in cell bodies to 15% of wild-type levels (Fig. 1D-E, Table S1). These results indicate that the PKD-2^C180S^ mutation prevents PKD-2 from both stabilizing into channel complexes and trafficking out of cell bodies, suggesting severe structural and trafficking defects caused by this single amino acid substitution.

The PKD-2^C180S^ mutation severely impaired sensory function to levels comparable to the *pkd-2 (sy606)* null allele (17, 25). Male response efficiency, which measures the male’s ability to respond to contact with hermaphrodite mating partners and initiate backing behavior. Compared with the 100% response efficiency of wild-type males (n = 60), PKD-2^C180S^ males showed a response efficiency of 20% (n = 60, P < 0.0001, Fig. 1F). Vulva location behavior, which assesses the ability to stop at the hermaphrodite vulva, was similarly impaired (Fig. 1G). While wild-type males stop at the vulva upon first encounter, both *pkd-2 (sy606)* null and PKD-2^C180S^ mutants passed the vulva and continued scanning (Fig. 1G; (17, 25). Thus, a single PKD-2^C180S^ substitution reduces protein levels, abolishes ciliary and EV trafficking, and eliminates sensory functions similar to null mutants.

### The PKD-2^C180S^ allele is recessive and does not interfere with wild-type PKD-2 localization

Cystogenesis in ADPKD can result from four potential genetic mechanisms: gain-of-function, dominant negative, haploinsufficiency, or molecular recessive loss-of-function (8). In a gain-of-function mechanism, the mutant protein acquires aberrant activity, rendering it constitutively active (27). In a dominant negative mechanism, the mutant protein interferes with the function of the wild-type protein (28). Haploinsufficiency arises when a single functional copy is insufficient to maintain normal cellular output (29). Finally, molecular-level recessive action occurs when the mutant allele is functionally silent in the presence of a wild-type copy yet drives disease upon loss of the second allele (30, 31). To directly test which mechanism underlies the PKD-2^C180S^ mutation, we leveraged the power of *C. elegans* genetics to visualize proteins produced from individual chromosomes simultaneously. We crossed a *pkd-2 (my122)* PKD-2::mScarlet (mSc) strain (24) with PKD-2::GFP or PKD-2^C180S^::GFP strains to generate F1 heterozygous males expressing dual-color fluorescent reporters (Fig. 2A-B).

**Figure 2.**
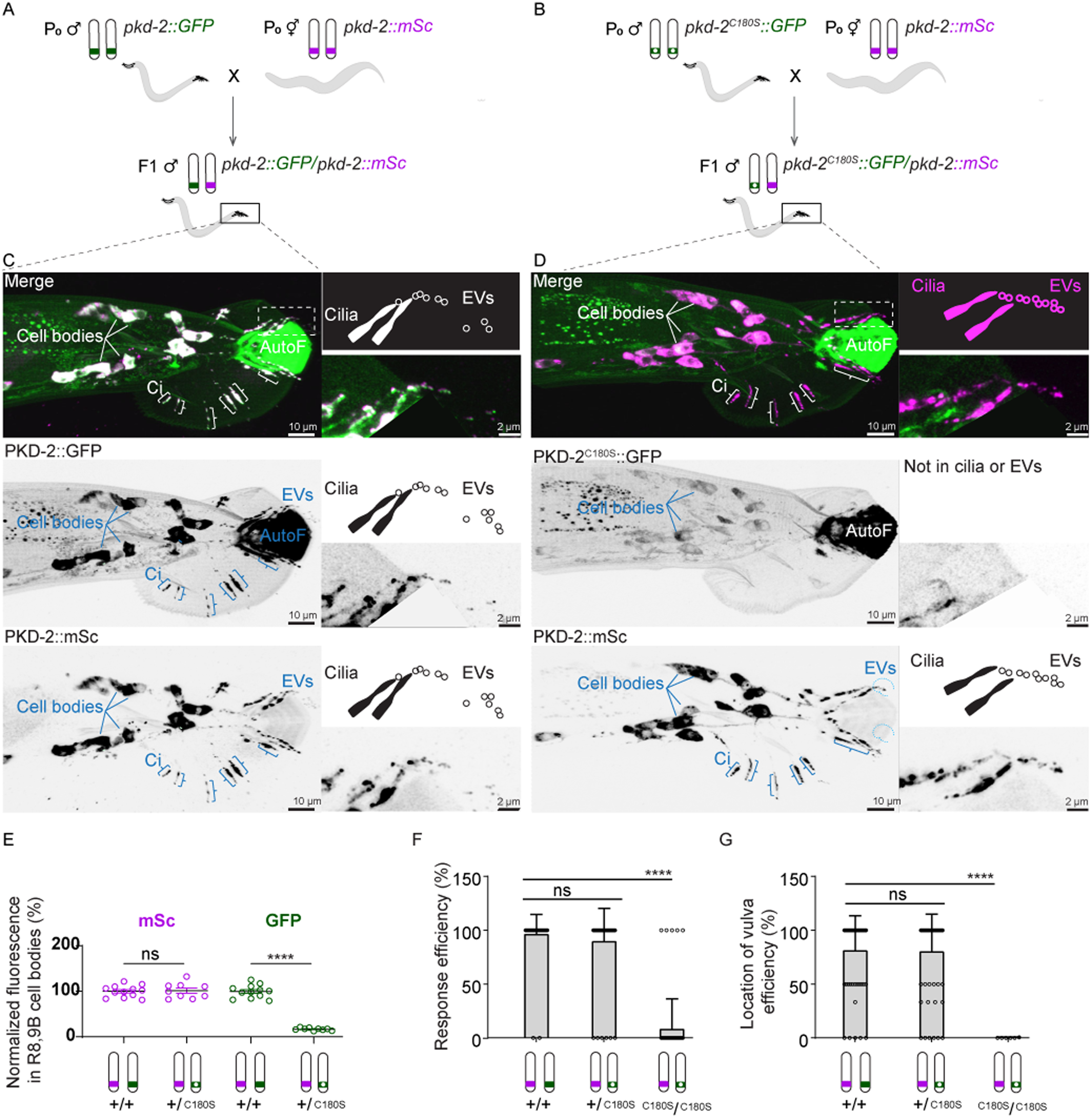
The PKD-2 ^C180S^ mutation was recessive in both cellular localization and sensory function. (A-B) Schematic diagram of experimental design: We tagged the wild-type copy of *pkd-2* with mScarlet (mSc) and crossed these hermaphrodites with males carrying either PKD-2::GFP (A) or PKD-2^C180S^::GFP (B). We examined mSc/GFP heterozygous, first generation (F1) male cross progeny and visualized proteins encoded from each chromosome simultaneously. For details, see Methods. (C-D) Representative micrographs of heterozygous male tails expressing PKD-2::GFP/PKD-2::mSc (C) and PKD-2^C180S^::GFP/PKD-2::mSc (D), respectively. The upper, middle, and lower panels show the merged, GFP, and mSc channels, respectively. The rectangle indicates the region shown in the insets to the right. AutoF denotes male tail autofluorescence, which is prominent in the GFP channel but reduced in the mSc channel. For insets, autofluorescence in the tail region is masked by background color to clearly display PKD-2 localization in cilia and EVs. Cartoons above each inset micrograph illustrate the corresponding cilia and EV structures. Cell bodies (lines), cilia (Ci, bracket), and EVs (dashed circles) are labeled. (C) Insets provide zoomed-in views of cilia and EVs, highlighting colocalization of PKD-2::GFP and PKD-2::mSc in these structures. A slight channel shift between GFP and mSc signals in the merged panel further confirms colocalization of PKD-2::GFP and PKD-2::mSc within EVs. (D) Insets provide zoomed-in views showing the absence of PKD-2^C180S^::GFP from cilia and EVs, while PKD-2::mSc remains localized to these structures. (E) Fluorescence intensity quantification of mSc and GFP in males with genotypes PKD-2::GFP/PKD-2::mSc and PKD-2^C180S^::GFP/PKD-2::mSc. Ray 8B and 9B neuronal cell bodies were used for fluorescence intensity quantification. Data were presented as mean ± SD. GFP and mSc fluorescence intensities were normalized to PKD-2::GFP/PKD-2::mSc, respectively. Statistical analysis was performed by one-way ANOVA with Bonferroni multiple comparisons (n = 9 and 12 for PKD-2::GFP/PKD-2::mSc and PKD-2^C180S^::GFP/PKD-2::mSc, respectively). **** indicates *p* < 0.0001; ns indicates *p* > 0.9999. (F-G) Mating behavior efficiency for response to hermaphrodite contact and vulva location behavior. Data were presented as mean ± SD. **** indicates *p* < 0.0001; n.s. indicates *p* > 0.9999. Statistical analysis was performed by Kruskal-Wallis test with Dunn’s multiple comparisons.

In PKD-2::GFP/PKD-2::mSc heterozygous males, both fluorescent proteins localized normally to the cell bodies, cilia, and ciliary EVs (Fig. 2C). However, in PKD-2^C180S^::GFP/PKD-2::mSc heterozygous males, only the wild-type protein trafficked normally (Fig. 2D). In these heterozygotes, PKD-2::mSc maintained wild-type localization patterns and levels, while PKD-2^C180S^::GFP remained restricted to the cell body at reduced levels, similar to homozygous PKD-2^C180S^ mutants (Fig. 2C-E, Fig. 1D). These results demonstrate that PKD-2^C180S^ acts in a recessive manner, with no dominant-negative effect on wild-type PKD-2 protein levels, trafficking to cilia, or localization to ciliary EVs.

### PKD-2^C180S^ does not interfere with wild-type PKD-2 function in male mating behavior

To assess whether PKD-2^C180S^ interferes with wild-type PKD-2 sensory function, we examined male mating behavior in heterozygotes. PKD-2^C180S^::GFP/PKD-2::mSc males exhibited normal mating behavior, performing indistinguishably from PKD-2::GFP/PKD-2::mSc heterozygotes (Fig. 2F-G). Consistent with the finding that the *pkd-2* null allele acts recessively and *pkd-2 (null)*/+ heterozygotes behave normally in mating behaviors (25), these behavioral results demonstrate that PKD-2 is haplosufficient in *C. elegans* and that PKD-2^C180S^ does not exert a dominant-negative effect on wild-type PKD-2 function. Together with the protein localization data (Fig. 2C-E), these findings establish that the PKD-2^C180S^ mutation acts recessively at both the cellular and organismal levels.

### PKD-2^C180S^ abolishes ciliary and EV localization of LOV-1

PC1 ciliary localization depends on PC2, mediated by coiled-coil interactions between their C-terminal cytosolic tails (32–35). Pathogenic PC2 TOP domain mutations (W414G, E442G) reduce PC1 ciliary localization (34, 36–39). In *C. elegans*, the *pkd-2* null mutant reduces LOV-1 cell body levels and abolishes LOV-1 ciliary localization (40). To determine whether the PKD-2^C180S^ variant impacts LOV-1 localization and cell body protein levels, we examined LOV-1 distribution in PKD-2^C180S^ mutant and compared the findings to those observed in *pkd-2* null animals.

In wild-type males, LOV-1 localizes to cell bodies, dendritic vesicles, cilia, and ciliary EVs (Fig. 3A; (24, 40). The PKD-2^C180S^ mutation abolished LOV-1::mSc ciliary and EV localization (Fig. 3B) and reduced LOV-1 levels in cell bodies to 11% of wild type (Fig. 3B, D), comparable to the *pkd-2 (sy606)* null allele (13%; Fig. 3C-D). Thus, PKD-2^C180S^ is functionally equivalent to a *pkd-2* null allele with respect to LOV-1 localization and stability.

**Figure 3.**
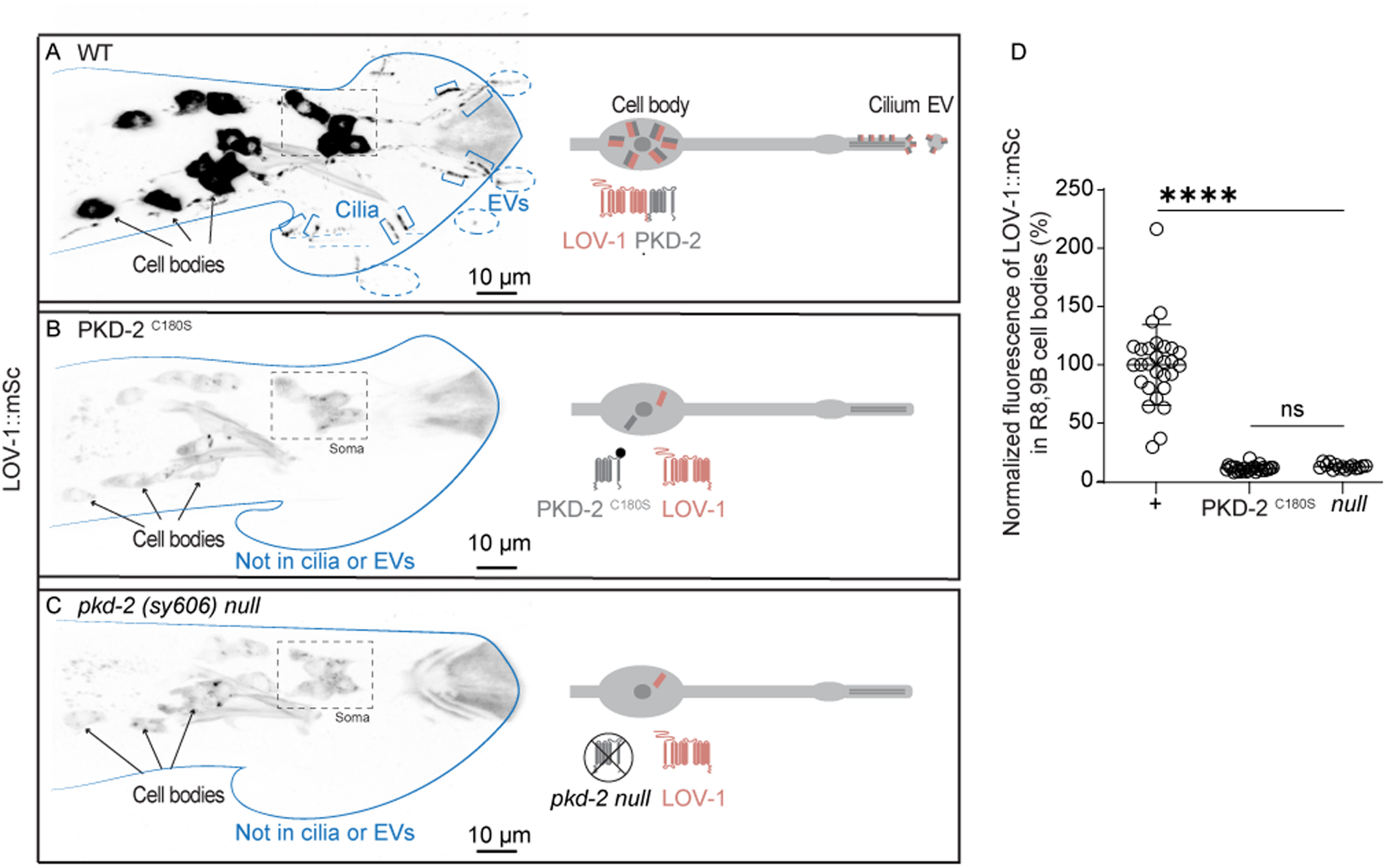
The PKD-2^C180S^ mutation decreased LOV-1 levels in the cell bodies and abolished LOV-1 localization to cilia and EVs. (A-C) Representative micrographs of LOV-1::mSc in wild-type (top), PKD-2^C180S^ (middle), and *pkd-2 (sy606)* null mutant (bottom) animals. Cartoons depict the localization and assembly status of the indicated proteins. Rectangles indicate the cell body regions used for fluorescence intensity quantification in panel D. (A) In wild-type animals, LOV-1::mSc was enriched in the cell bodies (arrows), sensory cilia (Ci, bracket), and EVs (dashed circles). We hypothesize that LOV-1::mSc (red) assembles with untagged PKD-2 (gray) to exit the cell bodies and localize to cilia and EVs. (B-C) In PKD-2^C180S^ and *pkd-2 (sy606)* null mutant animals, LOV-1::mSc levels were reduced in the cell bodies (arrows) and LOV-1::mSc was absent from cilia and EVs. We hypothesize that LOV-1::mSc (red) cannot assemble with PKD-2^C180S^ (B) and cannot exit the cell body in the absence of wild-type PKD-2 (C). (D) Quantification of LOV-1::mSc fluorescence intensity in the cell bodies of ray B neurons (rays 8 and 9). Data represent mean ± SD. Statistical significance was determined by the Kruskal-Wallis test with Dunn’s multiple comparisons. ns, *p* > 0.9999; ****, *p* < 0.0001.

These data demonstrate that the C180 residue is required for LOV-1 ciliary trafficking, likely through the essential role of C180 in disulfide bond formation and stabilization of inter-subunit interactions within the PKD-2•LOV-1 complex (Fig. 1B, Fig. 3E). The loss of LOV-1 ciliary localization in PKD-2^C180S^ mutants indicates that LOV-1 may traffic to cilia as part of a LOV-1•PKD-2 complex. Moreover, the severe reduction in cell body LOV-1 levels supports the established role of PC2 as a molecular chaperone for PC1 maturation and stability in the ER(11, 34).

### Reduced PKD-2^C180S^ in cell bodies is independent of LOV-1

To investigate how LOV-1 regulates PKD-2 localization dynamics in relation to PKD-2^C180S^, we generated a new null allele of *lov-1*, *lov-1 (my90)*, by inserting a STOP cassette into the second exon of the *lov-1* gene after Leucine 26 (Fig. S2A; for details, see Methods). *lov-1 (my90)* reduced PKD-2 levels in cell bodies to 60% of wild-type levels (Fig. S2B–D, K), an effect indistinguishable from that of the previously characterized genetic null allele *lov-1 (sy582)*.

These genetic results suggest that PKD-2 can assemble into homomers in the absence of LOV-1. In *+/*PKD-2^C180S^ heterozygotes, PKD-2^C180S^ does not interfere with wild-type PKD-2 cell body protein levels, ciliary trafficking, or EV trafficking, suggesting that PKD-2^C180S^ does not assemble into complexes with either PKD-2 homomers or LOV-1•PKD-2 heteromers. To test these two possibilities, we examined PKD-2^C180S^ protein levels in *lov-1 (my90)* mutants. PKD-2^C180S^::GFP cell body levels were indistinguishable between wild-type (16.21% ± 3.41%, mean ± SD) and *lov-1 (my90)* backgrounds (15.05% ± 5.31%, mean ± SD; Fig. 4A–D), demonstrating that in the absence of LOV-1, PKD-2^C180S^ is neither stabilized nor further destabilized.

**Figure 4.**
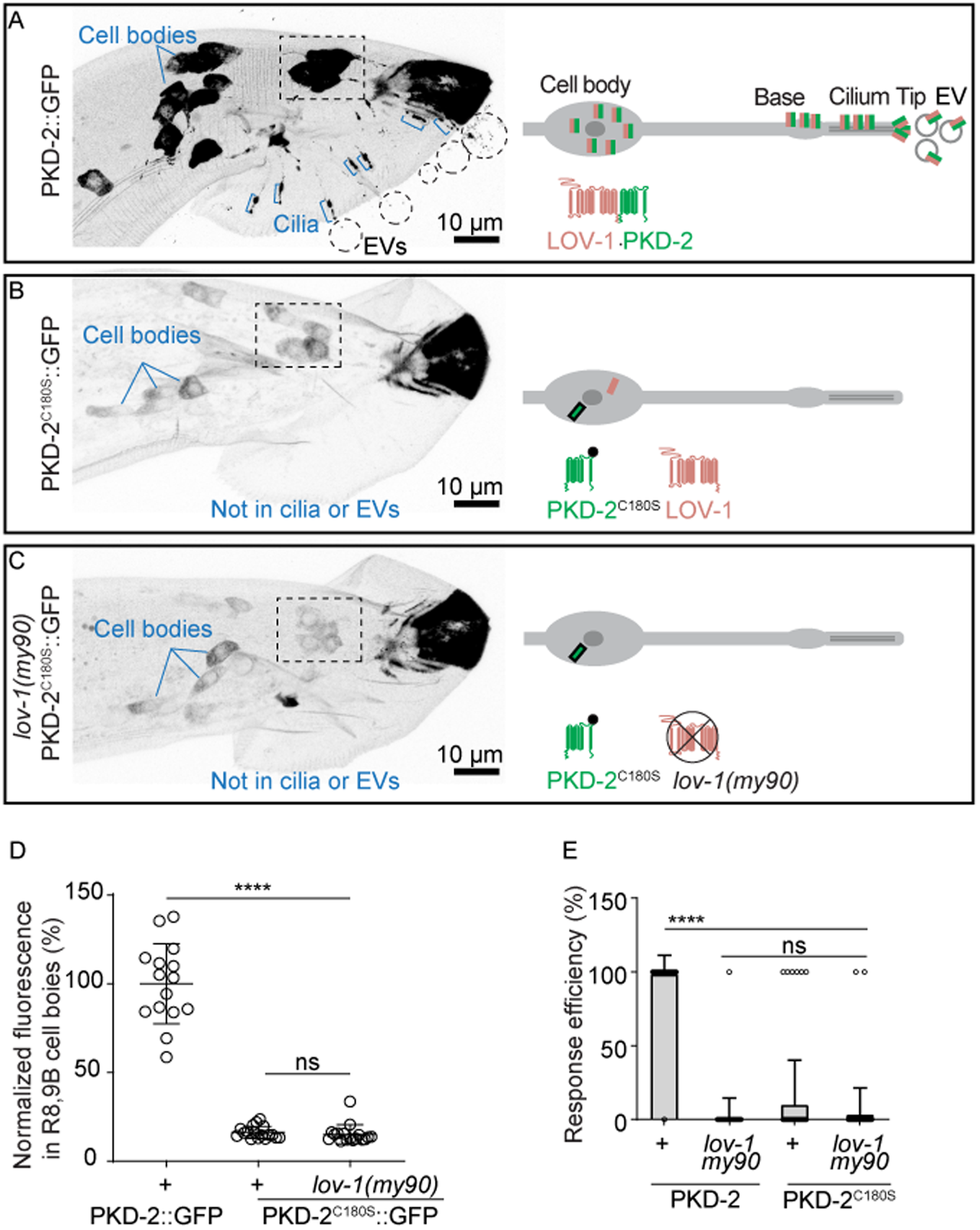
Decreased PKD-2^C180S^ protein levels in the cell body were independent of LOV-1. (A-C) Representative micrographs of animals expressing PKD-2::GFP (A), PKD-2^C180S^::GFP (B), and PKD-2^C180S^::GFP in a *lov-1(my90)* null mutant background (C). Rectangles indicate ray 8B and 9B cell bodies used for fluorescence intensity quantification in panel D. (A) In wild-type animals, PKD-2::GFP was enriched in the cell bodies (lines), sensory cilia (Ci, bracket), and EVs (dashed circles). We hypothesize that PKD-2::GFP assembles with untagged LOV-1 to exit the cell body and localize to cilia and EVs. (B–C) In PKD-2^C180S^ and *lov-1 (my90),* PKD-2^C180S^ mutant animals, PKD-2^C180S^ levels were reduced in the cell bodies (lines), and PKD-2^C180S^ was absent from cilia and EVs. We hypothesize that PKD-2^C180S^ cannot assemble with LOV-1 (B–C) or with itself (C), and cannot exit the cell body. (D) Quantification of PKD-2::GFP and PKD-2^C180S^::GFP fluorescence intensity in the cell bodies of ray B neurons (rays 8 and 9). Genotypes: PKD-2::GFP in wild-type, PKD-2^C180S^::GFP in wild-type, and PKD-2^C180S^::GFP in *lov-1 (my90)* mutant backgrounds (*n* = 15, 17, and 17, respectively). (E) Mating efficiency of wild-type, *lov-1 (my90)*, PKD-2^C180S^::GFP, and *lov-1(my90),* PKD-2^C180S^::GFP animals. (D–E) Data are presented as mean ± SD. Statistical analysis was performed using the Kruskal-Wallis test with Dunn’s multiple comparisons correction. n.s., *p* > 0.9999; ****, *p* < 0.0001.

Consistent with the cellular localization phenotype, male mating behavior was also defective in *lov-1 (my90)* mutants, as assessed by the response efficiency of males upon contact with hermaphrodite mating partners. Notably, the response efficiency was not significantly different between PKD-2^C180S^ single mutants and *lov-1 (my90);* PKD-2^C180S^ double mutants.

Taken together, our results demonstrate that in heterozygotes, PKD-2^C180S^ does not interfere with wild-type PKD-2 trafficking to cilia and EVs (Fig. 2). Furthermore, PKD-2^C180S^ mutant animals exhibited loss of LOV-1 localization to cilia and EVs, with reduced LOV-1 cell body levels comparable to those of *pkd-2* null mutants (Fig. 3). These genetic findings suggest that PKD-2^C180S^ acts at a step prior to LOV-1•PKD-2 heterocomplex assembly or PKD-2 homomer assembly.

### PKD-2 exhibits different dynamics on ciliary membranes with or without LOV-1

PC2 can exist as homotetramers or as heterotetramers in complex with PC1 (19–23). Although the PC1•PC2 complex has different ion channel properties compared to PC2 homotetramers (7, 41), the in vivo physiological significance of PC1•PC2 complexes versus PC2 homotetramers remains unknown. We therefore examined the ciliary and EV localization dynamics of PKD-2 in the presence or absence of LOV-1. PKD-2 can localize to cilia and EVs in the absence of LOV-1 (24, 26, 40), suggesting that PKD-2 can traffic independently. To directly compare the localization dynamics of putative LOV-1•PKD-2 heteromeric complexes versus putative PKD-2 homomers in their native genomic context, we quantified PKD-2::GFP distribution in wild-type animals (where PKD-2 forms complexes with LOV-1) and *lov-1* mutants (where only PKD-2 is present).

We measured PKD-2::GFP levels across four subcellular compartments: cell body, ciliary base, ciliary shaft, and ciliary tip. In *lov-1* mutants, PKD-2::GFP levels were reduced in all compartments compared to wild type, but the degree of reduction varied by location (Fig. S2B-L, Table S1). Cell body and ciliary tip levels were reduced to 50∼60% of wild type, while ciliary base and shaft levels were more severely reduced to 20∼30% of wild type (Fig. S2B–L). Additionally, the number of PKD-2::GFP-positive EVs in *lov-1* mutants decreased to approximately 30% of wild type (Fig. S2I, Table S1). The more pronounced reduction of PKD-2 at the ciliary base and shaft compared to the cell body and ciliary tip in *lov-1* mutants suggests that LOV-1•PKD-2 complexes are more stable than PKD-2 alone on the ciliary base and shaft membranes. Furthermore, although ciliary tip levels were reduced to only 55% of wild type in *lov-1* mutants, the number of PKD-2-positive EV were reduced to 30% of wild type, indicating that LOV-1•PKD-2 complexes are more efficiently packaged into ciliary EVs than PKD-2 alone.

Overall, our in vivo *C. elegans* system supports a model whereby the dynamic localization of polycystins depends on complex assembly between LOV-1 and PKD-2. The PC2^C331S^ orthologous mutation may similarly affect complex assembly, disrupting both PC1•PC2 heteromeric complex formation and PC2 homomer assembly (Illustrated in Fig. 4 Cartoon).

## Discussion

By introducing the *C. elegans* orthologous mutation PKD-2^C180S^ via endogenous genome editing, we demonstrate that the likely pathogenic PC2^C331S^ variant is a recessive loss-of-function allele that severely reduces protein stability, eliminates sensory function, and abolishes ciliary and EV localization of both PKD-2 and the PC1 homolog LOV-1. Rather than interfering with wild-type PKD-2, the C180S mutation acts at a step prior to channel complex assembly and exerts no dominant-negative effects, suggesting that PC2^C331S^ may contribute to cystogenesis through loss of polycystin complex function. Together, these findings validate a generalizable *C. elegans* pipeline for the functional classification of polycystin variants and provide mechanistic insight into how disruption of the PC1•PC2 signaling unit contributes to ADPKD.

### PKD-2^C180S^ is a recessive loss-of-function allele

A fundamental question in ADPKD genetics is the mechanism by which a given variant drives disease — gain-of-function, dominant negative, haploinsufficiency, or recessive loss-of-function — each of which carries distinct implications for how dysfunction of the PC1•PC2 ciliary signaling complex ultimately leads to cystogenesis (8). Yet direct distinction among these mechanisms through in vivo observation of proteins produced from two alleles simultaneously has been lacking. Here, using dual-color fluorescent reporters in heterozygous *C. elegans*, we demonstrate that PKD-2^C180S^ acts recessively at both the cellular and organismal levels. In heterozygotes, wild-type PKD-2 traffics normally to cilia and EVs, maintains normal protein levels, and supports normal male mating behavior, with no detectable interference from the mutant allele. These results establish that PKD-2 is haplosufficient in *C. elegans*, consistent with the previously reported recessive nature of the *pkd-2* null allele (25), and that PKD-2^C180S^ functions as a loss-of-function allele that phenocopies the *pkd-2* null.

The recessive nature of PKD-2^C180S^ has important implications for understanding ADPKD pathogenesis. Because the PKD-2^C180S^ mutant protein cannot incorporate into polycystin complexes, PKD-2^C180S^ is effectively invisible to wild-type PKD-2 in heterozygous cells. This molecular-level recessive loss-of-function is consistent with the two-hit model of cystogenesis, in which somatic loss of the remaining wild-type allele is required to initiate cyst formation (8). Our findings therefore predict that PC2^C331S^ patients would require a second somatic hit to abolish polycystin function in renal epithelial cells, with potential implications for prognosis and the timing of therapeutic intervention.

### PC2^C331S^ disrupts an early step in polycystin biogenesis

A central finding of this study is that the PKD-2^C180S^ mutation acts prior to channel complex assembly. Several lines of evidence support this conclusion. First, PKD-2^C180S^ protein levels in the cell body are reduced to ∼15% of wild type, indicating that the mutant protein is largely degraded before it can exit the endoplasmic reticulum. Second, PKD-2^C180S^ cell body levels are indistinguishable in wild-type and *lov-1* mutant backgrounds, demonstrating that the defect is independent of LOV-1 and therefore occurs before LOV-1•PKD-2 complex formation. Third, PKD-2^C180S^ fails to interact with wild-type PKD-2 in heterozygous animals. Together, these results place the C180S molecular defects before complex assembly.

In the human PC2 structure, C331 forms a disulfide bond with C444 within the TOP domain, constraining the Finger 1 extension that mediates inter-subunit contacts critical for tetramer stability (19, 20, 22, 42). Loss of this disulfide bond likely causes misfolding of the TOP domain, triggering ER quality control and subsequent degradation. This mechanism is consistent with a growing body of evidence that TOP domain integrity is a prerequisite for polycystin maturation and ciliary trafficking: the pathogenic mutation PC2^R322Q^ is targeted for endoplasmic reticulum-associated degradation (ERAD) in HEK293 cells (43), and PC2^W414G^ and PC2^E442G^ disrupt ciliary localization in HEK293 cells, in mouse models, and in mouse-derived primary cell cultures (37, 44, 45)Together with our findings, these studies suggest that TOP domain mutations represent a class of variants that are recognized and eliminated by ER quality control before the polycystin complex can assemble (46).

The identity of the ER surveillance machinery that may recognize misfolded PKD-2^C180S^ remains to be determined. One candidate is the protein disulfide isomerase PDI-6, the *C. elegans* homolog of PDIA6, which is highly expressed in the male sensory neurons that express PKD-2 (47). PDIA6 forms a complex with DNAJB11 (48), mutations in which cause atypical polycystic kidney disease (49). This raises the possibility that PKD-2^C180S^ (PC2^C331S^) and DNAJB11-associated atypical ADPKD may share an overlapping mechanism centered on defective polycystin maturation in the ER, and that this quality control step may represent a therapeutically targetable point in the polycystin biogenesis pathway.

One important caveat is that ciliary localization outcomes for polycystin variants may depend on both expression level and cellular context. PC2 TOP domain mutations including PC2^C331S^, PC2^R322Q/W^, and PC2^R325P/Q^ retain ciliary localization when overexpressed in HEK293 cells (50), yet PC2^R322Q^ is targeted for ERAD in the same cell type (43). Overexpression of PC2 also induces an extensive unfolded protein response and alters ER morphology (51) raising the possibility that supraphysiological expression saturates the quality control machinery that would otherwise degrade misfolded variants. These considerations highlight the value of studying polycystin variants at endogenous expression levels in native cellular contexts, as achieved in our *C. elegans* CRISPR knock-in model.

### Differential dynamics of PKD-2 with and without LOV-1 on ciliary membranes

PC2 preferentially assembles into heterocomplexes with PC1 when coexpressed in *Xenopus* oocytes (52). However, PC2 can localize to cilia independently of PC1 (53, 54), indicating that PC2 homomers also localize to ciliary membranes. Although PC1•PC2 heterocomplexes exhibit distinct ion conductance properties compared to PC2 homomers (41, 52), our understanding of the differential physiological functions and subciliary localization of these two complex types remains limited. Here, we show that in *C. elegans*, the endogenous dynamics of LOV-1•PKD-2 complexes differ from those of PKD-2 alone: LOV-1•PKD-2 complexes are preferentially retained or stabilized at ciliary base and shaft membranes and are more efficiently packaged into ciliary EVs than PKD-2 alone (Table S1; model in FigS2. E-G).

### *C. elegans* as a platform for testing ADPKD variants

Our study demonstrates the utility of *C. elegans* for functional analysis of polycystin missense variants of uncertain significance and/or that are categorized likely pathogenic. CRISPR/Cas9 editing at the endogenous *pkd-2* locus enables study of variants in their native genomic context, avoiding overexpression artifacts, while real-time in vivo imaging of polycystin localization to cilia and ciliary EVs provides dynamic information currently unattainable in fixed-cell assays. The genetic tractability of *C. elegans* further allows rapid dominance testing, epistasis analysis, and quantitative behavioral readouts. Conservation of key structural features, including the TOP domain disulfide bond network, validates *C. elegans* as a relevant model for human variants in both PC2 and PC1. The workflow herein provides a generalizable pipeline for evaluating variants of unknown significance in ADPKD.

## Materials and Methods

### *C. elegans* Strains and Maintenance

*C. elegans* strains were maintained at 20°C on nematode growth medium (NGM) plates seeded with *Escherichia coli* OP50 using standard methods (57). The following alleles were used in this study: *pkd-2 (my121 PKD-2::GFP, pkd-2 (my122 PKD-2::mSc;* (24), *pkd-2 (my154;* this work), *pkd-2 (sy606, null reference allele;*(17), and *lov-1 (my90;* this work). WormBase was used throughout experimental planning and interpretation of results (58).

### Structural Modeling

Structural predictions and protein-protein interactions were generated using AlphaFold server (59). Images were generated by UCSF Chimera (60). The *C. elegans* PKD-2 structure (UniProt: Q9U1S7, residues 60-619) was compared to the human PC2 structure (residues 185-723, PDB: 6A70) in complex with PC1. AlphaFold was also used to predict interactions between the TOP domains of three CePKD-2 subunits and LOV-1, with particular attention to disulfide bond formation at the interface of adjacent subunits.

### CRISPR/Cas9 Genome Editing

The *pkd-2 (my154)* allele was generated in the *my121* (24) background by introducing a point mutation changing cysteine 180 to serine using CRISPR/Cas9-mediated base editing. *lov-1 (my90)* was generated by inserting a STOP cassette (61) into the second exon of the *lov-1* gene after Leucine 26. For *lov-1 (my90),* the gRNA sequence 5’-GGGAA GUUUG UCCAG AGCAG-3’ and single-stranded oligonucleotide (ssODN) repair template 5’-ACTTT TAAGA TACAA ATTGA TGGAT TGCAT TACCA ACTGG GAAGT TTGTC CAGAG CAGAG GTGAC TAAGT GATAA GGATC CTCTT GACGG GATCG CAACT TTTCG ATTAG ACAAC GATGA CACAA CAATT GGAG-3’ were used. The edit was confirmed by PCR amplification using primers *lov-1 (my90)*F (5’-CCGCC TTTTC GCTTT TCGAC-3’) and *lov-1 (my90)*R (5’-CAGGT GAGTA ACGCC GACAT-3’). For *pkd-2 (my154),* the gRNA sequence 5’-AUCGU UUGCU UGGGG AACCU-3’ and ssODN repair template 5’-GGACG GAAAC TTCCA ATTCG ACGGA TAACG AGAAT ATGAT CTACT ATGAGA ATCGTT TGCTT GGGGA ACCTC GAATC AGAAT GTTGA AAGTG ACAAA TGACT CGTCT ACTGT GATGA AAAGT TTCCA GCGGG AGATT AAGGA ATGTT TTGCA AATTA TGAGG AAAAG CTCGA GGATA AGACG ATGGT C-3’ were used. The edit was confirmed by PCR amplification using primers *pkd-2 (my154)*F (5’-GTAAT GAGCG ACCTG TTTGT GG-3’) and *pkd-2 (my154)R* (5’-GGCTA ATCGC TGGAT CGACA G-3’). All genome edits were confirmed by DNA sequencing.

### Male Mating Behavior Assays

Response to hermaphrodite and vulva location assays were performed as previously described (17) with minor modifications. Briefly, a mating assay lawn was prepared by placing 10 µL of *E. coli* OP50 culture grown in LB broth onto an NGM plate and allowing it to dry with the lid open in a 37°C incubator for 3 to 4 hours to form a circular lawn approximately 0.5 cm in diameter. Ten minutes prior to each assay, 10–12 *unc-31* young adult hermaphrodites were transferred onto the mating lawn and allowed to acclimate. Male mating behavior was scored as follows: 1–3 males of the test genotype were released at the center of the lawn and observed for 5 minutes. A positive response was scored when a male contacted a hermaphrodite with its tail and initiated backward scanning along the hermaphrodite’s body. Vulva location efficiency was calculated as the reciprocal of the number of vulva contacts made by the male before stopping at the vulva. Prior to each assay, CB1490: *him-5 (e1490)V* males were used as positive (wild-type) controls, and PT9: *pkd-2 (sy606) IV; him-5 (e1490)V* males were used as negative controls to ensure normal mating behavior. Three independent blinded behavioral assays were performed, with investigators observing 20 males per genotype per assay (n = 60 total animals per genotype). Investigators scoring behavior were blinded to genotype.

### Airyscan super-resolution microscopy

Super-resolution imaging was performed on the Zeiss LSM880 confocal system equipped with Airyscan Super-Resolution Detector with 7 Single Photon Lasers (405, 458.488, 514, 561, 594, 633nm), Axio Observer 7 Motorized Inverted Microscope, Motorized X-Y Stage with Z-Piezo, T-PMT. MBS and Filter sets for GFP (excitation 488 nm) and mSc (excitation 561 nm) were used. Adult male animals were mounted on 10% agarose pads and immobilized using 10 mM levamisole. Images were acquired using consistent exposure times and laser power settings across all genotypes within each experiment. For localization studies, z-stack images were collected at 0.5 μm intervals through the entire volume of neurons of interest. Maximum intensity projections or single focal planes are presented as indicated in figure legends.

For quantitative imaging, ray B neurons (rays 4, 8 and 9) of the nine pairs of rays in the male tail were imaged. Sensory cilia were identified by their characteristic morphology and position. Extracellular vesicles (EVs) were identified as discrete puncta outside the cilia in the male tail region. At least three independent imaging sessions were performed for each experiment.

### Generation of F1 Male *C. elegans* Animals

To generate F1 males with specific PKD-2 allelic combinations, we used the temperature-sensitive *pha-1 (e2123ts) III* mutation as a genetic selection marker. Hermaphrodites carrying *pha-1 (e2123ts) III; pkd-2 (my122[PKD-2::mSc]) IV; him-5(e1490)* were crossed with *him-5* males expressing either PKD-2::GFP (control) or PKD-2^C180S^::GFP (test). Cross plates were maintained at 22°C, a restrictive temperature at which only cross-progeny (carrying the wild-type *pha-1(+)* allele from the male) could survive, while self-fertilized progeny (homozygous for *pha-1 (e2123ts*)) would arrest. We confirmed the effectiveness of this strategy by verifying that all surviving F1 males carried both fluorescent markers (Fig. 2D-E).

This same genetic selection approach was used for generating F1 males for both imaging experiments and male mating behavior assays. For all experiments, L4-stage F1 males were isolated 24 hours prior to analysis and maintained at 20°C to ensure developmental synchronization. Control strains underwent identical handling procedures to maintain consistent experimental conditions across all genotypes.

### Profiling of PKD-2::GFP distribution along cilia and at the ciliary base

Fluorescence quantification of the ciliary compartment was based on (62). Z stack acquisition for each male tail was performed to encompass environmental EVs, the cilium, and the ciliary base. Images were captured using a Plan-Apochromat 63x/1.40 Oil objective with a 2x zoom acquisition area under the Airyscan super-resolution mode. Raw image files were processed using the Airyscan Processing program within the Zen Black 2.0 software. Maximum projection images of each z stack were analyzed using the Profiling Program in Zen Blue software. The mean fluorescence intensity and the distance from the ciliary tip were recorded in an Excel file for subsequent data analysis. The fluorescence intensity of the distal cilium was determined by measuring 0 – 1000 nm from the ciliary tip, delineating the end of PKD-2 ciliary tip enrichment. The fluorescence intensity of the proximal cilium was determined by a 1000-2000 nm distance from the ciliary tip, whereas the ciliary base fluorescence intensity was determined from 2000 nm from the ciliary tip to the end of the PKD-2::GFP signal at the ciliary base. Relative fluorescence intensity was normalized to the mean fluorescence intensity of the distal ciliary PKD-2::GFP.

### Image Analysis and Quantification

Images were saved and processed using Zen Black software (Zeiss) to generate maximum intensity projections, and quantification was performed using the Zen Blue software (Zeiss) image analysis wizard. For cell body fluorescence intensity measurements, all images within the same experimental set were thresholded using identical parameters. For ray 8B and 9B neuronal cell body quantification in F1 males with genotypes *pkd-2::gfp/pkd-2::mSc* and *pkd-2^C180S^::gfp/pkd-2::mSc*, all images were thresholded using the mSc channel, and mean fluorescence intensities from both GFP and mSc channels were recorded in Microsoft Excel for further analysis.

### Statistical Analysis

The Prism software package (GraphPad Software 8) was used to carry out statistical analyses. Information about statistical tests, p values and n numbers are provided in the respective figures and figure legends.

## Supporting information

Wang_2026_Movie_S1

## Acknowledgements

This work was supported by National Institutes of Health (NIH) DK059418 and DK116606 (M.M.B), the Polycystic Kidney Disease Foundation Award #959686 (J.D.W. and J. W.), and #1443775 (I.A.N.). We thank Gloria Androwski for excellent technical assistance, Barr lab members and the Rutgers C. elegans community for feedback and constructive criticism throughout this project. We also thank WormBase, Japan National Bioresource Project for the nematode and Caenorhabditis Genetics Center (CGC) for resources information and strains. The CGC is supported by the National Institutes of Health - Office of Research Infrastructure Programs (P40OD010440).

---

**Figure S1.**
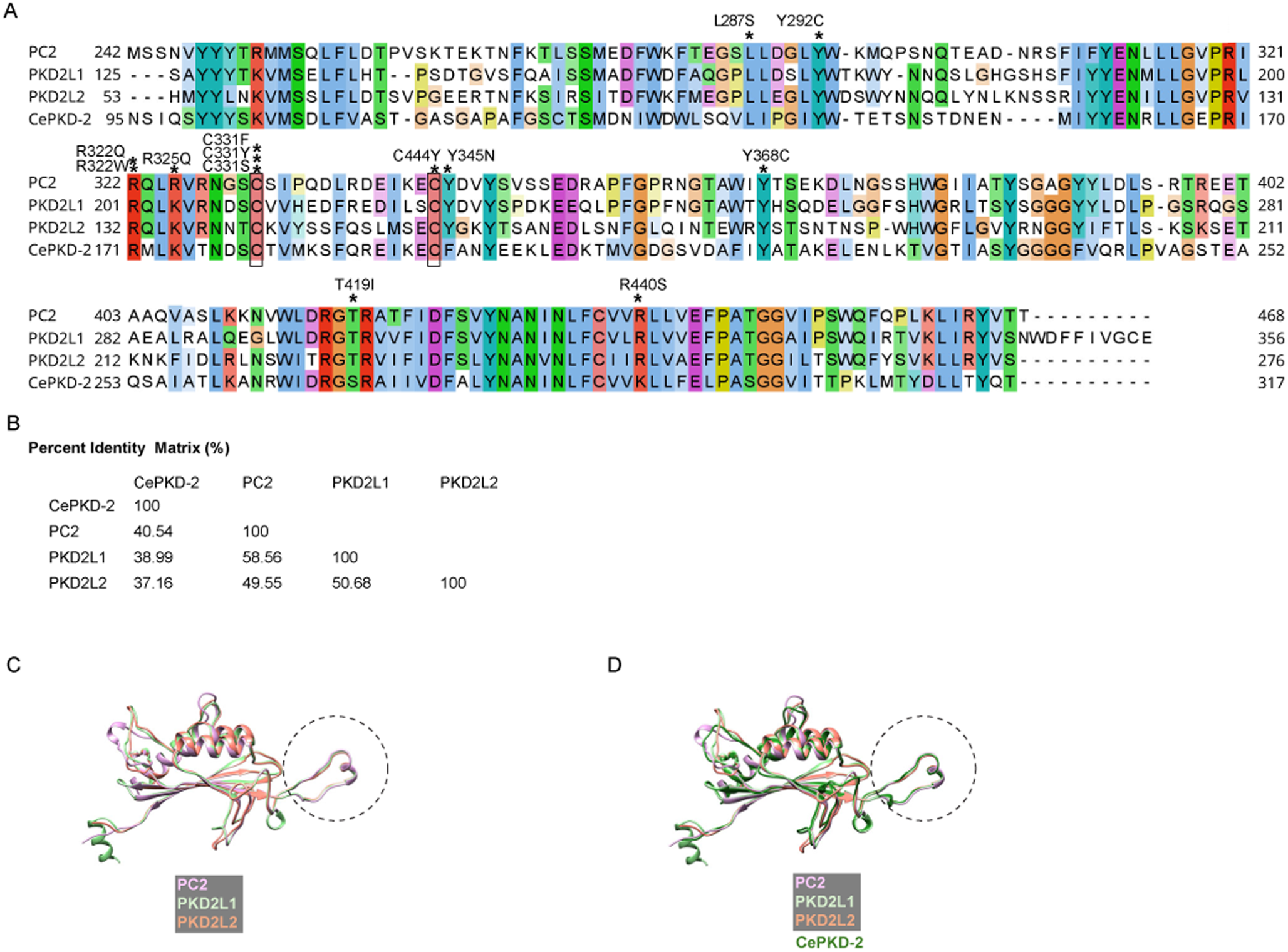
TOP domains were evolutionarily conserved in primary sequence and structure. (A) Multiple sequence alignment of TOP domains from human polycystin-2 family members (PC2, PKD2L1, and PKD2L2) and *C. elegans* PKD-2 (CePKD-2). Sequences were aligned using MUSCLE (55) and visualized with Jalview (56). Pathogenic and likely pathogenic missense variants documented in ClinVar are marked with asterisks above the corresponding residues, with variant names annotated alongside. Rectangles indicate cysteine residues forming disulfide bonds in the Finger 1 structure of the TOP domain. Protein sequences are listed as protein name, residue range, and UniProt ID: PC2, 242–468, Q13563; PKD2L1, 125–356, Q9P0L9; PKD2L2, 53–276, Q9NZM6; CePKD-2, 95–317, Q9U1S7. (B) Percent identity matrix among TOP domains of CePKD-2, PC2, PKD2L1, and PKD2L2. CePKD-2 was most similar to PC2 (40.54%), followed by PKD2L1 (38.99%) and PKD2L2 (37.16%). (C) Structural comparison of TOP domains from PC2 family members (predicted by AlphaFold Server), showing aligned core α-helices, β-sheets, and Finger 1 regions (circled). (D) AlphaFold-predicted CePKD-2 TOP domain structure aligned with PC2 family members.

**Figure S2.**
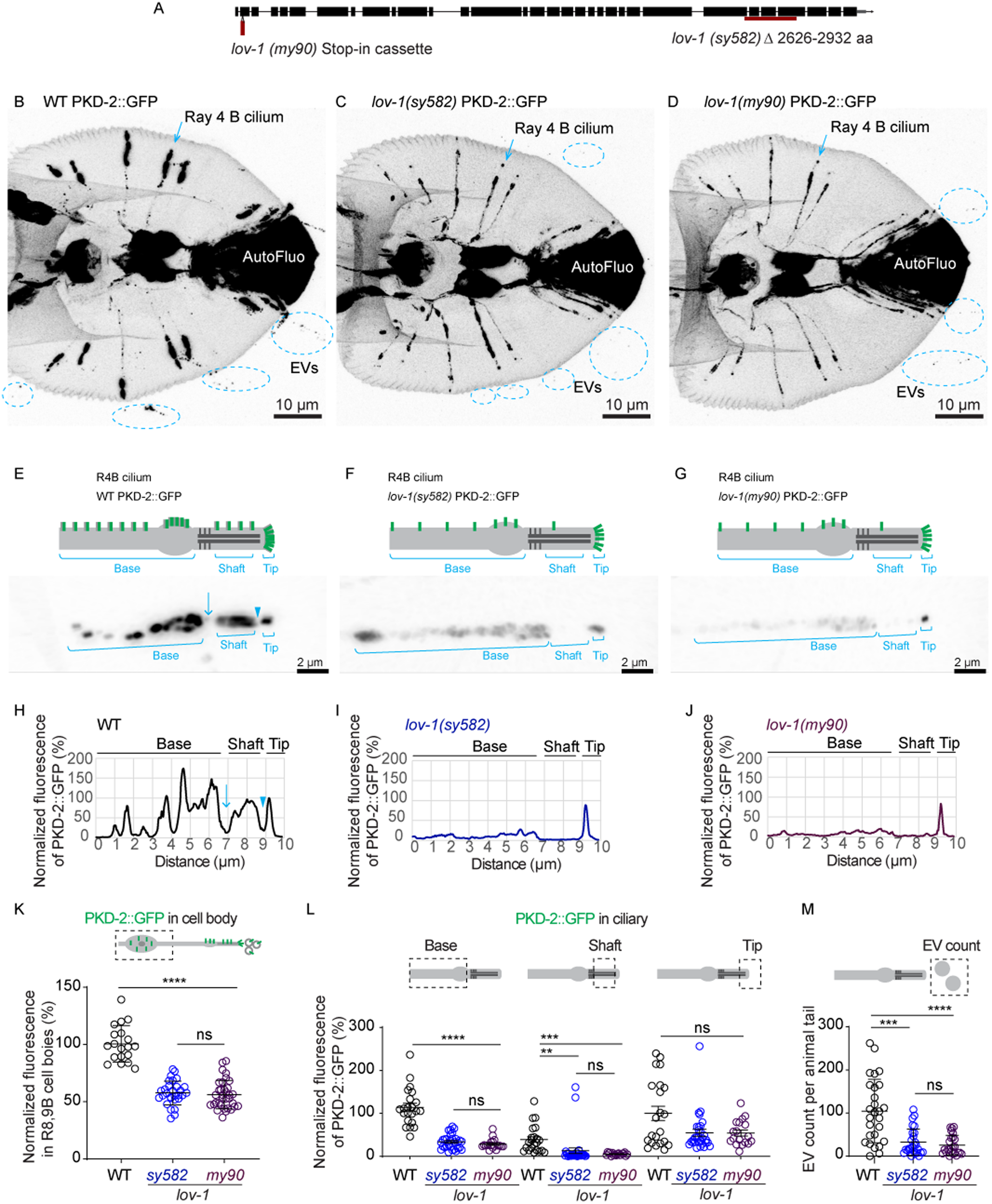
PKD-2 with and without LOV-1 displayed distinct dynamic subcellular localization. (A) Schematic of the *lov-1* gene structure and two *lov-1* mutant alleles. The *lov-1 (my90)* allele was a null allele generated by inserting a stop-in cassette into the second exon, resulting in protein truncation at leucine 26 (see Methods). The *lov-1 (sy582)* allele carries an in-frame deletion of residues 2626–2932 and is a genetic null (17, 25). (B–D) Representative micrographs of PKD-2::GFP in wild-type (B), *lov-1(sy582)* (C), and *lov-1 (my90)* (D) mutant male tails. In *lov-1* mutants, PKD-2::GFP was visible in the ciliary tip and base, while the shaft signal of PKD-2::GFP was reduced (C–D). Arrows indicate the ray 4B neuron cilium used for fluorescence profiling in panels E–G. (E–G) PKD-2::GFP fluorescence profiles of representative ray 4B cilia from panels B–D, respectively. In (E), an open arrow indicates a gap in PKD-2::GFP signal between the base (periciliary membrane compartment, PCMC) and the ciliary shaft, and an arrowhead indicates a gap between the shaft and the tip. In *lov-1* mutants, PKD-2 levels were lowest in the shaft region between the two gaps, as shown in the aligned fluorescence profiles (H–J). (H–J) Fluorescence profiles of ray 4B cilia in wild-type (H), *lov-1 (sy582)* (I), and *lov-1 (my90)* (J) animals, respectively. (K) Quantification of PKD-2::GFP fluorescence intensity in the cell bodies of ray 8B and 9B neurons in wild-type, *lov-1 (sy582)*, and *lov-1 (my90)* mutant animals (*n* = 19, 29, and 34, respectively). (L) Quantification of PKD-2::GFP fluorescence intensity at the ciliary base, shaft, and tip of ray 4B neurons in wild-type, *lov-1 (sy582)*, and *lov-1 (my90)* mutant animals (*n* = 22, 30, and 18 cilia, respectively). (M) Number of PKD-2::GFP-positive EVs in the tails of wild-type, *lov-1 (sy582)*, and *lov-1 (my90)* mutant male animals. Each dot represents the EV count from a single animal (*n* = 26, 25, and 24 animals, respectively). (K–M) Data are presented as mean ± SD. Statistical analysis was performed using the Kruskal-Wallis test with Dunn’s multiple comparisons correction. n.s., *p* > 0.9999; **, *p* < 0.01; ***, *p* < 0.001; ****, *p* < 0.0001.

**Table S1.**
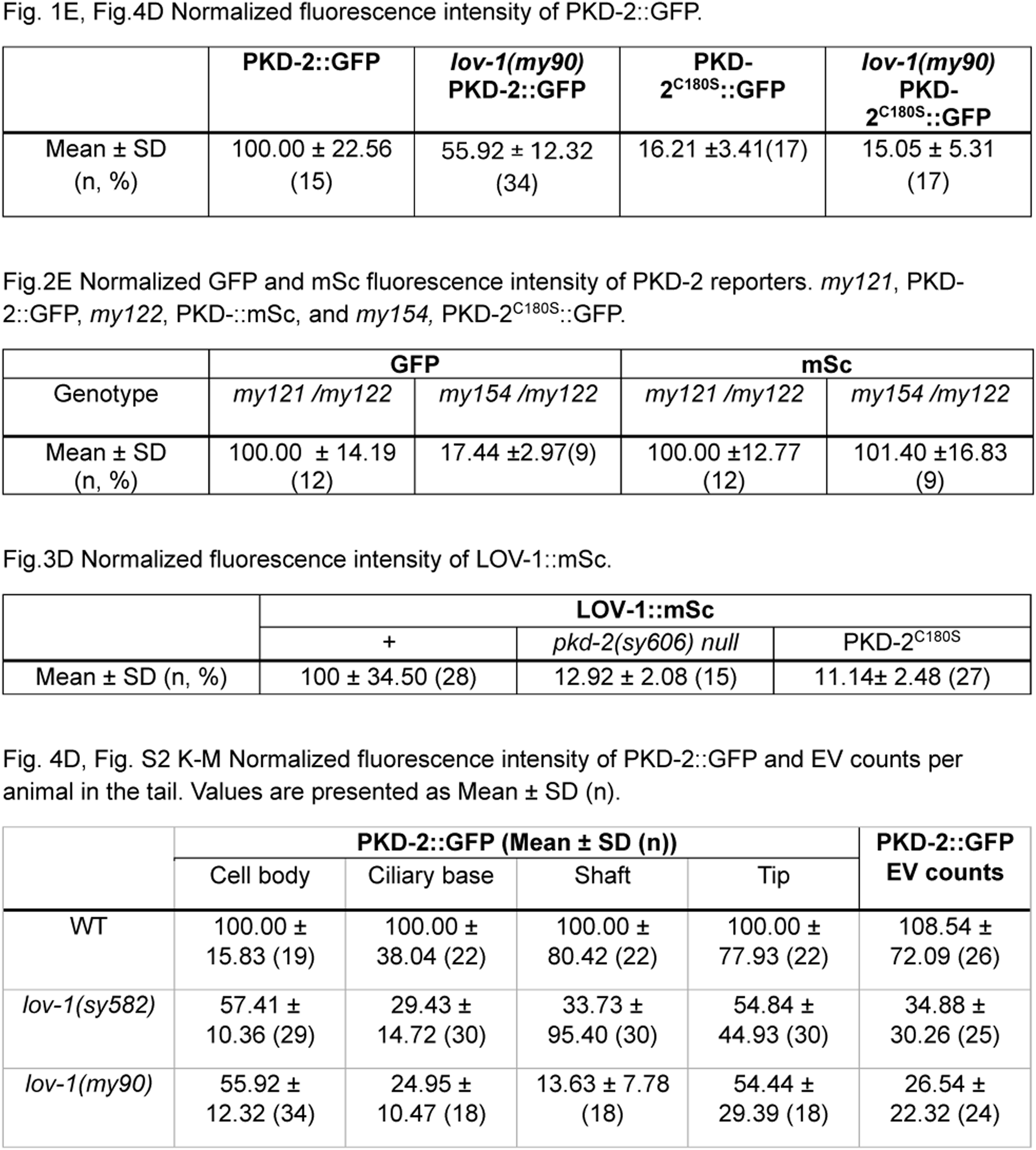
Source data for fluorescence intensity quantification graphs in this study.

**Table S2.** strains used this study.

| *C. elegans* Strain | genotype | Reference |
| --- | --- | --- |
| CB1490 | *him-5* (e1490) *V* | (60) |
| CB169 | *unc-31* (e169) *IV* | (55) |
| PT3775 | *pkd-2* (*my121*[*pkd-2::gfp*] *IV*; *him-5* *V* | (24) |
| PT3783 | *pkd-2* (*my122*[*pkd-2::mSc*] *IV*; *him-5* (e1490) *V* | (24) |
| PT4280 | *pkd-2* (*my154* [*PKD-2*<sup>C180S</sup>::*GFP*] *IV*; *him-5* (e1490) *V* | This work |
| PT4075 | *pha-1* (e2123) *III*; *pkd-2* (*my122*[*pkd-2::mSc*] *IV*; *him-5*(e1490) *V* | This work |
| PT9 | *pkd-2* (*sy606*) *IV*; *him-5*(e1490) *V* | (17) |
| PT3629 | *lov-1* (*my78*[*lov-1::mSc*] *II*; *him-5*(e1490) *V* | (40) |
| PT4354 | *lov-1* (*my78* [*LOV-1::mSc*] *II*; *pkd-2* (*sy606*) *IV*; *him-5*(e1490) *V* | This work |
| PT4032 | *lov-1* (*sy582*) *II*; *pkd-2* (*my121*[*pkd-2::gfp*] *IV*; *him-5* (e1490) *V* | This work |
| PT4060 | *lov-1* (*my90*[*lov-1*(*STOP*)] *II*; *pkd-2* (*my121*[*pkd-2::gfp*] *IV*; *him-5*(e1490) *V* | This work |
| PT4162 | *lov-1* (*my90*) *II*; *pkd-2* (*my154* [*PKD-2*<sup>C180S</sup>::*GFP*] *IV*; *him-5* (e1490) *V* | This work |

## Video S1

Structural comparison of AlphaFold-predicted structures of CePKD-2 (blue, UniProt: Q9U1S7, residues 60–619) and human PC2 (white, residues 185–723, PDB: 6A70). CePKD-2 cysteine 180 and human PC2 cysteine 331 are highlighted in purple. Video was generated by UCSF Chimera.

## References

1. Yamamoto S, Kanca O, Wangler MF & Bellen HJ (2023) Integrating non-mammalian model organisms in the diagnosis of rare genetic diseases in humans. Nature Reviews Genetics 25(1): 46–60.

2. Chebib FT, Hanna C, Harris PC, Torres VE & Dahl NK (2025) Autosomal dominant polycystic kidney disease: A review. JAMA 333(19): 1708–1719.

3. Cornec-Le Gall E, Alam A & Perrone RD (2019) Autosomal dominant polycystic kidney disease. The Lancet 393(10174): 919–935.

4. Hughes J, et al (1995) The polycystic kidney disease 1 (PKD1) gene encodes a novel protein with multiple cell recognition domains. Nat Genet 10(2): 151–160.

5. Maser RL, Calvet JP & Parnell SC (2022) The GPCR properties of polycystin-1-A new paradigm. Frontiers in Molecular Biosciences 9

6. Mochizuki T, et al (1996) PKD2, a gene for polycystic kidney disease that encodes an integral membrane protein. Science 272(5266): 1339–1342.

7. Douguet D, Patel A & Honoré E (2019) Structure and function of polycystins: Insights into polycystic kidney disease. Nature Reviews Nephrology 15(7): 412–422.

8. Qiu J, Germino GG & Menezes LF (2023) Mechanisms of cyst development in polycystic kidney disease. Advances in Kidney Disease and Health 30(3): 209–219.

9. Hilgendorf KI, Myers BR & Reiter JF (2024) Emerging mechanistic understanding of cilia function in cellular signalling. Nature Reviews Molecular Cell Biology 25(7): 555–573.

10. Boletta A & Caplan MJ (2025) Physiologic mechanisms underlying polycystic kidney disease. Physiol Rev 105(3): 1553–1607.

11. Hu J & Harris PC (2020) Regulation of polycystin expression, maturation and trafficking. Cell Signal 72

12. Pisitkun T, Shen R & Knepper MA (2004) Identification and proteomic profiling of exosomes in human urine. Proceedings of the National Academy of Sciences 101(36): 13368–13373.

13. Hogan MC, et al (2009) Characterization of PKD protein-positive exosome-like vesicles. Journal of the American Society of Nephrology 20(2): 278–288.

14. Hogan MC, et al (2015) Identification of biomarkers for PKD1 using urinary exosomes. Journal of the American Society of Nephrology 26(7): 1661–1670.

15. Lea WA, et al (2020) Analysis of the polycystin complex (PCC) in human urinary exosome–like vesicles (ELVs). Scientific Reports 10(1)

16. Hogan MC & Ward CJ (2024) An extracellular vesicle based hypothesis for the genesis of the polycystic kidney diseases. Extracellular Vesicle 4: 100048.

17. Barr MM & Sternberg PW (1999) A polycystic kidney-disease gene homologue required for male mating behaviour in C. elegans. Nature 401(6751): 386–389.

18. Wang J, et al (2014) C. elegans ciliated sensory neurons release extracellular vesicles that function in animal communication. Current Biology 24(5): 519–525.

19. Su Q, et al (2018) Structure of the human PKD1-PKD2 complex. Science 361(6406)

20. Shen PS, et al (2016) The structure of the polycystic kidney disease channel PKD2 in lipid nanodiscs. Cell 167(3): 763–773.

21. Grieben M, et al (2017) Structure of the polycystic kidney disease TRP channel polycystin-2 (PC2). Nat Struct Mol Biol 24(2): 114–122.

22. Wilkes M, et al (2017) Molecular insights into lipid-assisted Ca2+ regulation of the TRP channel polycystin-2. Nat Struct Mol Biol 24(2): 123–130.

23. Wang Q, et al (2020) Lipid interactions of a ciliary membrane TRP channel: Simulation and structural studies of polycystin-2. Structure 28(2): 169–184.

24. Nikonorova IA, et al (2025) Polycystins recruit cargo to distinct ciliary extracellular vesicle subtypes in C. elegans. Nature Communications 16(1)

25. Barr MM, et al (2001) The caenorhabditis elegans autosomal dominant polycystic kidney disease gene homologs lov-1 and pkd-2 act in the same pathway. Current Biology 11(17): 1341–1346.

26. Bae YK, et al (2006) General and cell-type specific mechanisms target TRPP2/PKD-2 to cilia. Development 133(19): 3859–3870.

27. Kurbegovic A, et al (2010) Pkd1 transgenic mice: Adult model of polycystic kidney disease with extrarenal and renal phenotypes. Hum Mol Genet 19(7): 1174–1189.

28. Gallagher AR, et al (2006) A truncated polycystin-2 protein causes polycystic kidney disease and retinal degeneration in transgenic rats. Journal of the American Society of Nephrology 17(10): 2719–2730.

29. Chang MY, et al (2006) Haploinsufficiency of Pkd2 is associated with increased tubular cell proliferation and interstitial fibrosis in two murine Pkd2 models. Nephrology Dialysis Transplantation 21(8): 2078–2084.

30. Tan AY, et al (2018) Somatic mutations in renal cyst epithelium in autosomal dominant polycystic kidney disease. Journal of the American Society of Nephrology 29(8): 2139–2156.

31. Zhang Z, et al (2021) Detection of PKD1 and PKD2 somatic variants in autosomal dominant polycystic kidney cyst epithelial cells by whole-genome sequencing. Journal of the American Society of Nephrology 32(12)

32. Casuscelli J, et al (2009) Analysis of the cytoplasmic interaction between polycystin-1 and polycystin-2. American Journal of Physiology - Renal Physiology 297(5)

33. Su X, et al (2015) Regulation of polycystin-1 ciliary trafficking by motifs at its C-terminus and polycystin-2 but not by cleavage at the GPS site. J Cell Sci 128(22): 4063–4073.

34. Gainullin VG, Hopp K, Ward CJ, Hommerding CJ & Harris PC (2015) Polycystin-1 maturation requires polycystin-2 in a dose-dependent manner. J Clin Invest 125(2): 607–620.

35. Chapin HC, Rajendran V & Caplan MJ (2010) Polycystin-1 surface localization is stimulated by polycystin-2 and cleavage at the g protein-coupled receptor proteolytic site. Mol Biol Cell 21(24): 4338–4348.

36. Walker RV, et al (2019) Ciliary exclusion of polycystin-2 promotes kidney cystogenesis in an autosomal dominant polycystic kidney disease model. Nature Communications 10(1)

37. Cai Y, et al (2014) Altered trafficking and stability of polycystins underlie polycystic kidney disease. J Clin Invest 124(12): 5129–5144.

38. Grimes DT, et al (2016) Genetic analysis reveals a hierarchy of interactions between polycystin-encoding genes and genes controlling cilia function during left-right determination. PLoS Genetics 12(6)

39. Itabashi T, et al (2025) Cholesterol ensures ciliary polycystin-2 localization to prevent polycystic kidney disease. Life Science Alliance 8(4)

40. Walsh JD, et al (2022) Tracking N- and C-termini of C. elegans polycystin-1 reveals their distinct targeting requirements and functions in cilia and extracellular vesicles. PLoS Genetics 18(12)

41. Wang Z, et al (2019) The ion channel function of polycystin-1 in the polycystin-1/polycystin-2 complex. EMBO Rep 20(11)

42. Grieben M, et al (2017) Structure of the polycystic kidney disease TRP channel polycystin-2 (PC2). Nature Structural and Molecular Biology 24(2): 114–122.

43. Guerriero CJ, et al (2025) Identification of polycystin 2 missense mutants targeted for endoplasmic reticulum-associated degradation. American Journal of Physiology-Cell Physiology 328(2): C483–C499.

44. Grimes DT, et al (2016) Genetic analysis reveals a hierarchy of interactions between polycystin-encoding genes and genes controlling cilia function during left-right determination. PLOS Genetics 12(6): e1006070.

45. Walker RV, et al (2019) Ciliary exclusion of polycystin-2 promotes kidney cystogenesis in an autosomal dominant polycystic kidney disease model. Nat Commun 10(1): 4072.

46. Salehi-Najafabadi Z, et al (2017) Extracellular loops are essential for the assembly and function of polycystin receptor-ion channel complexes. J Biol Chem 292(10): 4210–4221.

47. Hammarlund M, Hobert O, Miller DM & Sestan N (2018) The CeNGEN project: The complete gene expression map of an entire nervous system. Neuron 99(3): 430–433.

48. Piette BL, et al (2021) Comprehensive interactome profiling of the human Hsp70 network highlights functional differentiation of J domains. Mol Cell 81(12): 2549–2565.

49. Busch T, et al (2024) The role of the co-chaperone DNAJB11 in polycystic kidney disease: Molecular mechanisms and cellular origin of cyst formation. FASEB Journal 38(21)

50. Vien TN, Wang J, Ng LCT, Cao E & DeCaen PG (2020) Molecular dysregulation of ciliary polycystin-2 channels caused by variants in the TOP domain. Proc Natl Acad Sci U S A 117(19): 10329–10338.

51. Heinz V, Rachel R & Ziegler C (2025) Application of STEM tomography to investigate smooth ER morphology under stress conditions. J Microsc

52. Staudner T, et al (2025) Ion channel function of polycystin-2/polycystin-1 heteromer revealed by structure-guided mutagenesis. FEBS Lett 599(12): 1649–1668.

53. Geng L, et al (2006) Polycystin-2 traffics to cilia independently of polycystin-1 by using an N-terminal RVxP motif. J Cell Sci 119(7): 1383–1395.

54. Liu X, et al (2018) Polycystin-2 is an essential ion channel subunit in the primary cilium of the renal collecting duct epithelium. Elife 7: e33183.

55. Madeira F, et al (2024) The EMBL-EBI job dispatcher sequence analysis tools framework in 2024. Nucleic Acids Res 52: W521–W525.

56. Waterhouse AM, Procter JB, Martin DMA, Clamp M & Barton GJ (2009) Jalview version 2--a multiple sequence alignment editor and analysis workbench. Bioinformatics 25(9): 1189–1191.

57. Brenner S (1974) The genetics of caenorhabditis elegans. Genetics 77(1): 71–94.

58. Sternberg PW, et al (2024) WormBase 2024: Status and transitioning to alliance infrastructure. Genetics 227(1)

59. Abramson J, et al (2024) Accurate structure prediction of biomolecular interactions with AlphaFold 3. Nature 630(8016): 493–500.

60. Pettersen EF, et al (2004) UCSF chimera--a visualization system for exploratory research and analysis. Journal of Computational Chemistry 25(13): 1605–1612.

61. Wang H, Park H, Liu J & Sternberg PW (2018) An efficient genome editing strategy to generate putative null mutants in caenorhabditis elegans using CRISPR/Cas9. G3 (Bethesda, Md.) 8(11): 3607–3616.

62. Wang J, et al (2024) Ciliary intrinsic mechanisms regulate dynamic ciliary extracellular vesicle release from sensory neurons. Current Biology 34(12): 2756–2763.e2.

